# Parasitophorous vacuole membranes of *Toxoplasma gondii* and *Plasmodium falciparum* lack the lipid asymmetry characteristic of host cell plasma membranes

**DOI:** 10.64898/2026.08.11.744334

**Authors:** Rikako Konishi, Yuri Nakashima, Tatsunori Masatani, Masahito Asada, Hakimi Hassan, Kayoko Fukuda, Sayuri Kuriyama, Yoshifumi Nishikawa, Osamu Kaneko, Vern B. Carruthers, Akikazu Fujita

## Abstract

Apicomplexan parasites, including *Toxoplasma gondii* and *Plasmodium falciparum*, reside within a specialized compartment known as the parasitophorous vacuole (PV) during their intracellular life cycle. The PV membrane (PVM), which derives from the host plasma membrane upon invasion, serves as a selective barrier that permits nutrient acquisition while shielding the parasite from host defense mechanisms. Although the protein composition of the PVM has been studied extensively, its lipid organization remains poorly understood. Using the quick-freeze, freeze-fracture replica labeling (QF-FRL) method, we quantitatively analyzed the transbilayer distribution of phosphatidylserine (PtdSer), phosphatidylethanolamine (PtdEtn), and GM3 ganglioside in the PVM of *T. gondii* and *P. falciparum*. Unlike host cell plasma membranes, where these lipids exhibit strict asymmetry—PtdSer and PtdEtn confined to the cytoplasmic leaflet and GM3 to the exoplasmic leaflet—we found that all three lipids were symmetrically distributed across both leaflets of the PVM. This striking loss of lipid asymmetry suggests that the PVM undergoes profound remodeling during infection. The presence of PtdSer and PtdEtn in the luminal leaflet may facilitate the binding of perforin-like proteins (PLP1s) during egress. These findings reveal a unique feature of the PVM that redefines our understanding of host–parasite membrane biology.

## 1. Introduction

Apicomplexan parasites are a large and diverse phylum of obligate intracellular protozoa that include major human and animal pathogens such as *Plasmodium falciparum*, the causative agent of malaria, and *Toxoplasma gondii*, the agent of toxoplasmosis. Malaria remains one of the world’s most devastating infectious diseases, with hundreds of millions of clinical cases each year and a particularly high mortality in young children living in tropical and subtropical regions [1, 2]. *T. gondii* infects nearly one-third of the global population, usually remaining latent but occasionally causing severe or life-threatening disease in immunocompromised individuals and during congenital infection [3]). Understanding how these parasites establish and maintain their intracellular niche is central to parasitology and infectious disease biology.

A defining feature of apicomplexan infection is the formation of the parasitophorous vacuole (PV), a membrane-bound compartment that houses the parasite during its intracellular stage. The PV membrane (PVM) originates from the host plasma membrane (PM) during invasion but undergoes extensive remodeling that alters its protein and lipid composition [4–6]. The PVM functions as a selective barrier, allowing nutrient exchange while protecting the parasite from cytoplasmic immune mechanisms. Host transmembrane proteins that anchored to the cytoskeleton are excluded at a structure known as the moving junction during invasion, whereas certain lipids and GPI-anchored proteins are incorporated, reflecting a selective process in membrane formation [4, 5, 7].

While considerable progress has been made in characterizing the protein components of the PVM, much less is known about its lipid architecture. Lipid asymmetry is a fundamental property of eukaryotic membranes, maintained by flippases and scramblases that regulate the distribution of phospholipids between the cytoplasmic (protoplasmic, PF) and exoplasmic (EF) leaflets. Phosphatidylserine (PtdSer) and phosphatidylethanolamine (PtdEtn) are typically confined to the PF, whereas glycosphingolipids such as GM3 are localized to the EF. This asymmetric organization is critical for cellular signaling, vesicle trafficking, and recognition of apoptotic or infected cells [8]. However, whether this lipid asymmetry is preserved or disrupted in the PVM remains unknown.

Previous approaches to studying membrane lipid topology have relied heavily on fluorescently tagged lipid-binding domains that can bind cytoplasmic lipid headgroups [9–11]. However, these probes are limited in their ability to resolve the luminal leaflet and can introduce artifacts by competing with endogenous lipid-binding proteins [12–14]. Electron microscopy (EM) can visualize membrane lipids in situ but typically lacks leaflet-specific resolution [15–17]. To overcome these limitations, we used the quick-freeze, freeze-fracture replica labeling (QF-FRL) technique, which allows high-resolution, leaflet-specific detection of lipid distribution [18].

In this study, we applied QF-FRL to examine the distribution of PtdSer, PtdEtn, and GM3 in both the host PM and the PVM of *T. gondii* and *P. falciparum*. Our analyses reveal a striking symmetry of these lipids across the two leaflets of the PVM, in contrast to the pronounced asymmetry of host PMs. This loss of asymmetry suggests extensive lipid rearrangement during PV formation, providing new insights into the molecular organization and potential functional properties of the PVM.

## 2. Materials and Methods

### 2.1. Probes

Complementary DNA generated by RT-PCR of total RNA of the mouse fibroblast and HEK293 cell were used as templates to clone of the C2 domain of MFGE8 [19] as previously described. A recombinant glutathione S-transferase (GST) fusion protein containing the MFG-E8 C2 domain (GST-MFGE8-C2) was expressed in *Escherichia coli* and purified as previously described [20, 21]. A secondary rabbit anti-GST antibody was purchased from Bethyl Laboratories (Montgomery, TX, USA). Anti-mouse and anti-rabbit IgG antibodies conjugated with 10 nm gold particles and an Alexa Fluor 488-conjugated anti-rabbit IgG antibody were purchased from Jackson ImmunoResearch Laboratories (West Grove, PA, USA). Biotin-conjugated duramycin-LC (biotin-duramycin) and mouse anti-GM3 and mouse anti-biotin antibodies were purchased from Polysciences (Warrington, PA, USA), SEIKAGAKU KOGYOU (Tokyo, Japan), and Jackson ImmunoResearch Laboratories, respectively.

### 2.2. Human foreskin fibroblast-1

Human foreskin fibroblast-1 (HFF-1) was cultured in Dulbecco’s modified Eagle’s medium (DMEM; GIBCO, Grand Island, NE, USA) supplemented with 10% fetal calf serum, 50 U/mL penicillin, and 0.05 mg/mL streptomycin at 37 °C under 5% CO_2_.

### 2.3. Parasite lines and culture

*T. gondii* tachyzoites of the PLK strain (kindly provided by Dr. Kami Kim, Albert Einstein College of Medicine, NY, USA) were maintained in HFF-1 cultured in Dulbecco’s modified Eagle’s medium (DMEM; GIBCO, Grand Island, NE, USA) supplemented with 10% fetal calf serum, 50 U/mL penicillin, and 0.05 mg/mL streptomycin at 37 °C under 5% CO_2_. For experiments, *T. gondii* tachyzoites were inoculated into confluent HFF-1 monolayers and incubated for 2–3 days post-infection before sample preparation. *P. falciparum* Dd2 parasite line was originally obtained from National Institute of Health, USA. The parasites were maintained with O^+^ human erythrocyte at 2% hematocrit in fibrinogen-free human plasma-containing complete RPMI1640 medium and transfection was performed as described [22]. Human erythrocytes and plasma were obtained from Japanese Red Cross Society with approval from the Research Ethics Committee, Institute of Tropical Medicine, Nagasaki University.

### 2.4. Fluorescence imaging of PtdSer

HFF-1 cells were plated on glass coverslips and, when necessary, infected with *T. gondii* tachyzoites. Two to three days post-infection, the cells were fixed with 4% paraformaldehyde in phosphate buffer (PB) and, if required, permeabilized with 0.05% saponin in phosphate-buffered saline (PBS). Samples were then incubated with GST-MFGE8-C2, followed by a rabbit anti-GST antibody (Bethyl Laboratories) and an Alexa Fluor 488-conjugated anti-rabbit IgG antibody, and examined under a fluorescence microscope (IX73; Olympus, Tokyo, Japan).

### 2.5. Quick-freezing and freeze-fracture

For quantitative lipid labeling of the biological membranes of *T. gondii* and *P. falciparum*, and HFF-1, the cells were quick-frozen by either a metal sandwich quick-freezing method or high-pressure freezing using an HPM 010 high-pressure freezing apparatus (Leica Microsystems, Wetzlar, Germany). For the metal sandwich freezing of *P. falciparum*-infected erythrocytes, a small volume of their pellets was placed on a copper foil, covered with a thin gold foil (∼4 mm^2^ area; 20 μm thickness), and then frozen using a quick press between two gold-plated copper blocks precooled in liquid nitrogen [21, 23]. For high-pressure freezing, *T. gondii*-infected HFF-1 cells cultured on the gold foil (∼4 mm^2^ area; 20 μm thickness) was sandwiched between flat aluminum discs (Engineering Office M. Wohlwend, Sennwald, Switzerland) and frozen according to the manufacturer’s instructions [24]. For quick-freezing of HHF-1, cells grown on a small gold foil (∼4 mm^2^ area; 20 μm thickness) were inverted on prewarmed 10% gelatin on a copper foil with the cell side down and processed according to the metal sandwich method described above [21, 23].

The frozen specimens were transferred to the cold stage of a Balzers BAF400 apparatus (Bal-Tec AG, Lichtenstein) and fractured at –130 °C under a vacuum of ∼1 × 10^−6^ mbar. Replicas were produced by electron-beam evaporation in three steps: carbon (C; ∼2 nm thickness) at an angle of 90°, platinum–carbon (Pt/C; 1–2 nm thickness) at an angle of 45°, and C (10–20 nm thickness) at an angle of 90° to the specimen surface (Fujita et al. 2010). The deposition thickness was adjusted using a crystal-thickness monitor (EM QSG100; Leica Microsystems).

The thawed specimens were treated overnight with 2.5% sodium dodecyl sulfate (SDS) in 0.1 M Tris-HCl (pH 8.0) at 60–70 °C, and the replicas were stored in 50% buffered glycerol at –30 °C until use.

### 2.6. Labeling and EM observation

Probe labeling was performed as previously described [18]. Briefly, after rinsing, the freeze-fracture replicas were blocked with PBS containing 2% cold-water fish-skin gelatin at room temperature for 30 min. The replicas were then incubated overnight at 4 °C with the anti-GM3 antibody, GST-MFGE8-C2 (73 ng/mL), or biotin-duramycin (60 μM) diluted in PBS containing 2% cold-water fish-skin gelatin. After four washes with PBS containing 0.1% bovine serum albumin (BSA), the replicas were incubated at 37 °C for 30 min with rabbit anti-GST antibody (5 μg/mL) or mouse anti-biotin antibody (130 μg/mL) for GST-MFGE8-C2 or biotin-duramycin, respectively, followed by incubation with 10 nm gold-conjugated anti-mouse IgM antibody, anti-rabbit, anti-mouse IgG antibody for anti-GM3 antibody, GST-MFGE8-C2, or biotin-duramycin, respectively, in PBS containing 2% cold-water fish-skin gelatin. The replicas were transferred to Formvar-coated grids and examined by transmission electron microscopy (TEM) (H7650; HITACHI, Tokyo, Japan) operated at 80 kV.

### 2.7. Statistical analysis

EM and fluorescence images from at least three independent experiments were used for the analyses. The number of colloidal gold particles was counted manually, and the areas were measured using ImageJ software [25]. Labeling density within each selected structure was calculated by dividing the number of colloidal gold particles by the corresponding area. For each structure, labeling density was measured in more than ten randomly captured micrographs. Fluorescence intensities of Alexa Fluor 488–labeled anti-rabbit IgG antibodies in HFF-1 cells were also quantified using ImageJ software. Statistical differences between the samples were analyzed using Student’s *t*-test.

## 3. Results

### 3-1. Overall distribution of PtdSer in uninfected and T. gondii-infected host cells

To determine whether *T. gondii* infection alters the overall cellular level or distribution of PtdSer, we first performed indirect immunofluorescence assay using probes GST-MFGE8-C2, rabbit anti-GST antibody, and Alexa Fluor 488–conjugated anti-rabbit IgG to detect PtdSer-specific signals in HFF-1 cells (Fig. 1). In uninfected HFF-1 cells, PtdSer labeling was detected mainly along the PF of the PM (Fig. 1A). In *T. gondii*-infected cells, PtdSer signals were also observed within the PF, but not EF, leaflet of the PM (Fig. 1B). Quantitative fluorescence intensity measurements using ImageJ showed no significant change in total PtdSer signal intensity between uninfected and infected cells (Fig. 1C), suggesting that infection does not alter the cellular abundance and cytoplasmic leaflet localization of PtdSer.

**Fig. 1.**
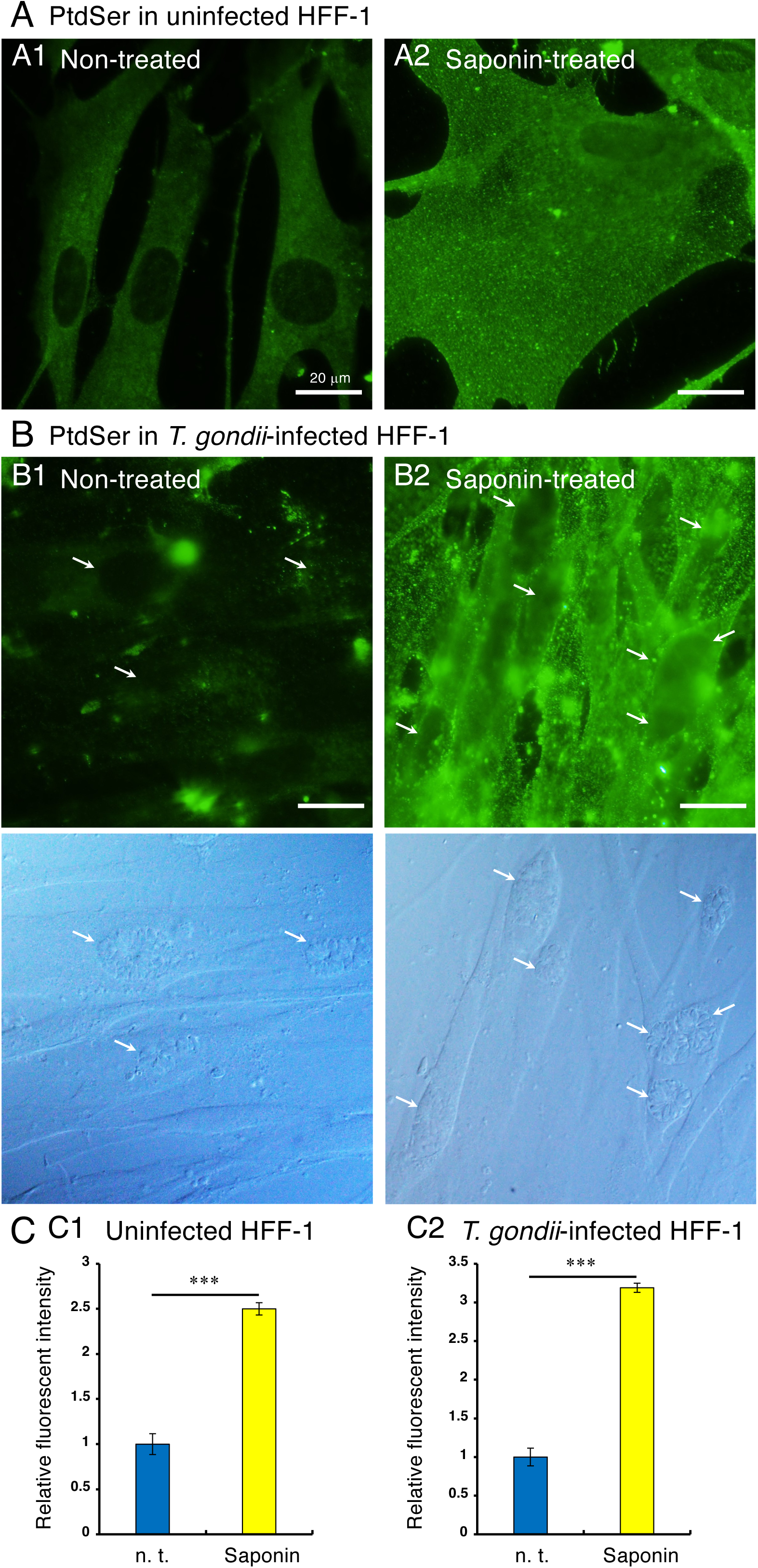
Overall distribution of PtdSer in uninfected and *T. gondii*–infected HFF-1 cells. Uninfected HFF-1 cells (A) and HFF-1 cells infected with *T. gondii* tachyzoites (B) were fixed with 4% paraformaldehyde, followed by treatment with 0.05% saponin (A2, B2) in PBS or left untreated (a). IFA images show PtdSer labeling using GST-MEFG8-C2, rabbit anti-GST-antibody, and Alexa Fluor 488–conjugated anti-rabbit IgG antibody. The lower panels in B display phase-contrast micrographs of the infected cells, where arrows indicate a rosette of intravacuolar tachyzoites. (C) The quantification of PtdSer labeling intensity in saponin-treated samples reveals significantly higher levels in the plasma membrane (yellow column) compared to untreated cells (blue column) in both uninfected and infected HFF-1 cells, indicating that PtdSer labeling is concentrated at the cytoplasmic, but not exoplasmic, sides of the plasma membrane in both the uninfected (A) and infected HFF-1 (B) cells. Scale bars: 20 μm. n. t.: Non-treated, *t*-test, ***, *p* < 0.001.

### 3-2. Leaflet-specific analysis of PtdSer in the host cell PM and PVM of T. gondii-infected cells

To visualize the leaflet-specific localization of PtdSer, we employed the QF-FRL technique. A schematic of the labeling procedure is shown in Fig. 2A. Briefly, cells were rapidly frozen, fractured to expose the EF and PF leaflets, and then the localization of colloidal gold-labeled PtdSer was examined by TEM. In replicas of uninfected HFF-1 cells (Fig. 2B), PtdSer labeling was almost exclusively observed on the PF (Fig. 2B2) of the PM, whereas the EF (Fig. 2B1) exhibited only background-level labeling. Gold particles were clearly distinguished from intramembrane particles (IMPs, yellow arrowheads), allowing unambiguous identification of labeled structures. In *T. gondii*-infected cells, both the EF and PF of the parasitophorous vacuole membrane (PVM) showed abundant PtdSer labeling (Fig. 2C), in sharp contrast to the asymmetric distribution observed in the host PM. Quantitative analysis confirmed that the PtdSer densities on the EF and PF of the PVM were nearly equivalent, whereas those in the PM were strongly biased toward the PF (Fig. 2D). These findings indicate that lipid asymmetry is lost during infection, leading to the exposure of PtdSer on both sides of the membrane.

**Fig. 2.**
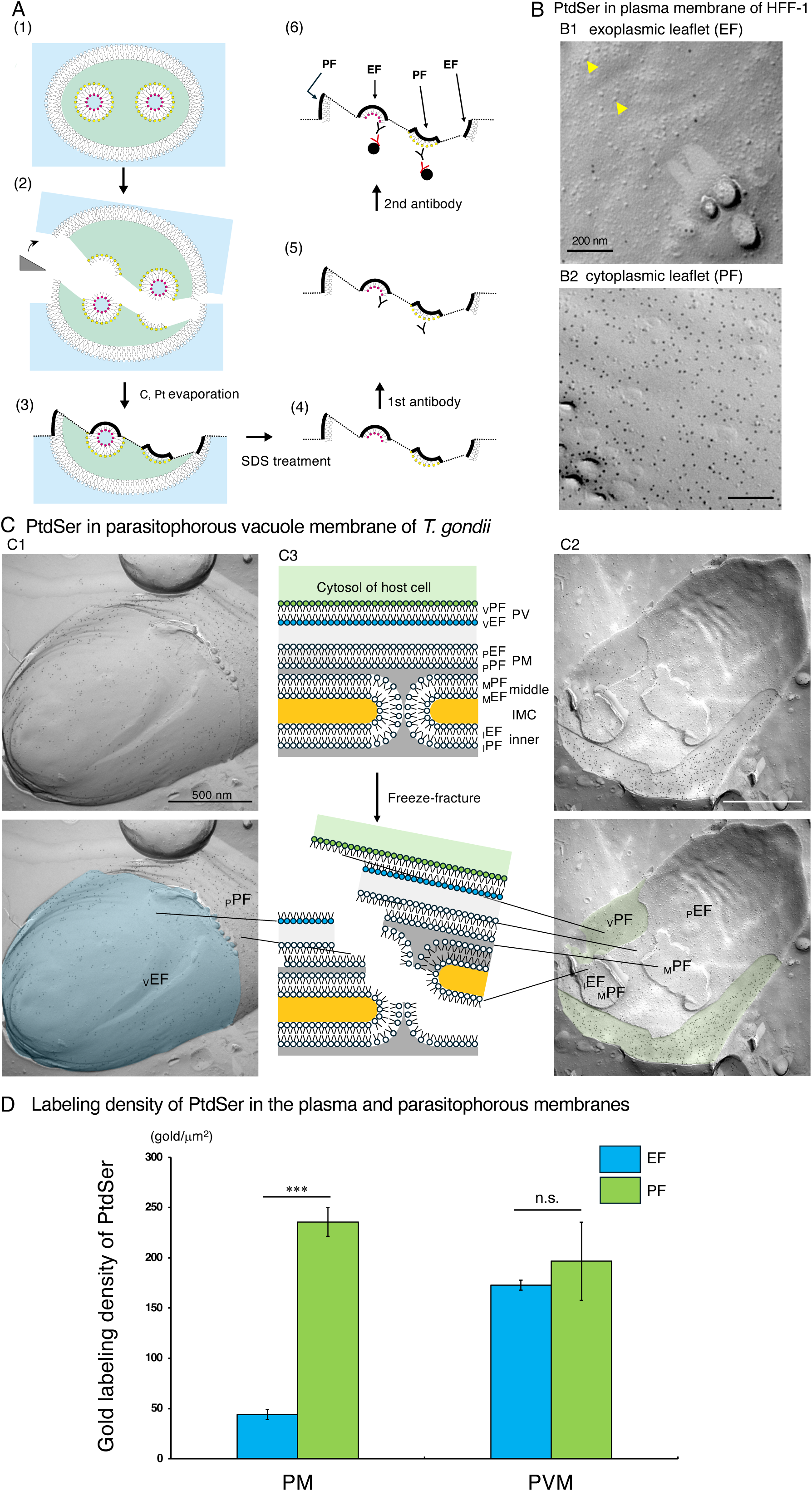
PtdSer distribution in the host plasma membrane and the parasitophorous vacuole membrane of *T. gondii*. (A) Schematic representation of the QF-FRL procedure showing labeling sites on the exoplasmic (EF) and protoplasmic (PF) leaflets. Fracture planes passing through the cytoplasm and the extracellular space are shown by a dotted line. (B) Representative replica image of the HFF-1 plasma membrane (PM) showing gold particles labeling of PtdSer (10-nm gold) and intramembrane particles (IMPs, yellow arrowheads). The image is annotated to distinguish labeled lipids from IMPs. Asymmetric localization of PtdSer in the EF (B1) and PF (B2) of the HFF-1 PM (PF > EF). (C) Replica image of the parasitophorous vacuole membrane (PVM) surrounding *T. gondii* tachyzoites, showing symmetric labeling of PtdSer across the EF (blue) and PF (green). The fracture faces of various membranes, including the parasitophorous vacuole (V), plasma membrane (P), middle IMC (inner membrane complex) membrane (M), and inner IMC membrane (I), are abbreviated as _V_PF, _V_EF, _P_PF, _P_EF, _M_PF, _M_EF, _I_PF, and _I_EF, respectively. (D) Quantitative analysis of PtdSer labeling densities in EF and PF of HFF-1 PM and *T. gondii* PVM. Values represent mean ± SE (n = 20 replicas). Scale bars: 200 nm (B), 500 nm (C). n. s.: not significant, *t*-test, ***, *p* < 0.001

### 3-3. Symmetrical localization of phosphatidylethanolamine in the PVM of T. gondii-infected cells

Next, we examined the distribution of PtdEtn, another aminophospholipid typically confined to the cytoplasmic leaflet of eukaryotic membranes. In uninfected HFF-1 cells, PtdEtn labeling was abundant on the PF but scarce on the EF of the PM (Fig. 3A). However, in *T. gondii*-infected cells, PtdEtn labeling was detected on both leaflets of the PVM (Fig. 3B). Quantitative evaluation revealed comparable labeling densities on the EF and PF of the PVM (Fig. 3C), again suggesting a breakdown of phospholipid asymmetry in the PVM.

**Fig. 3.**
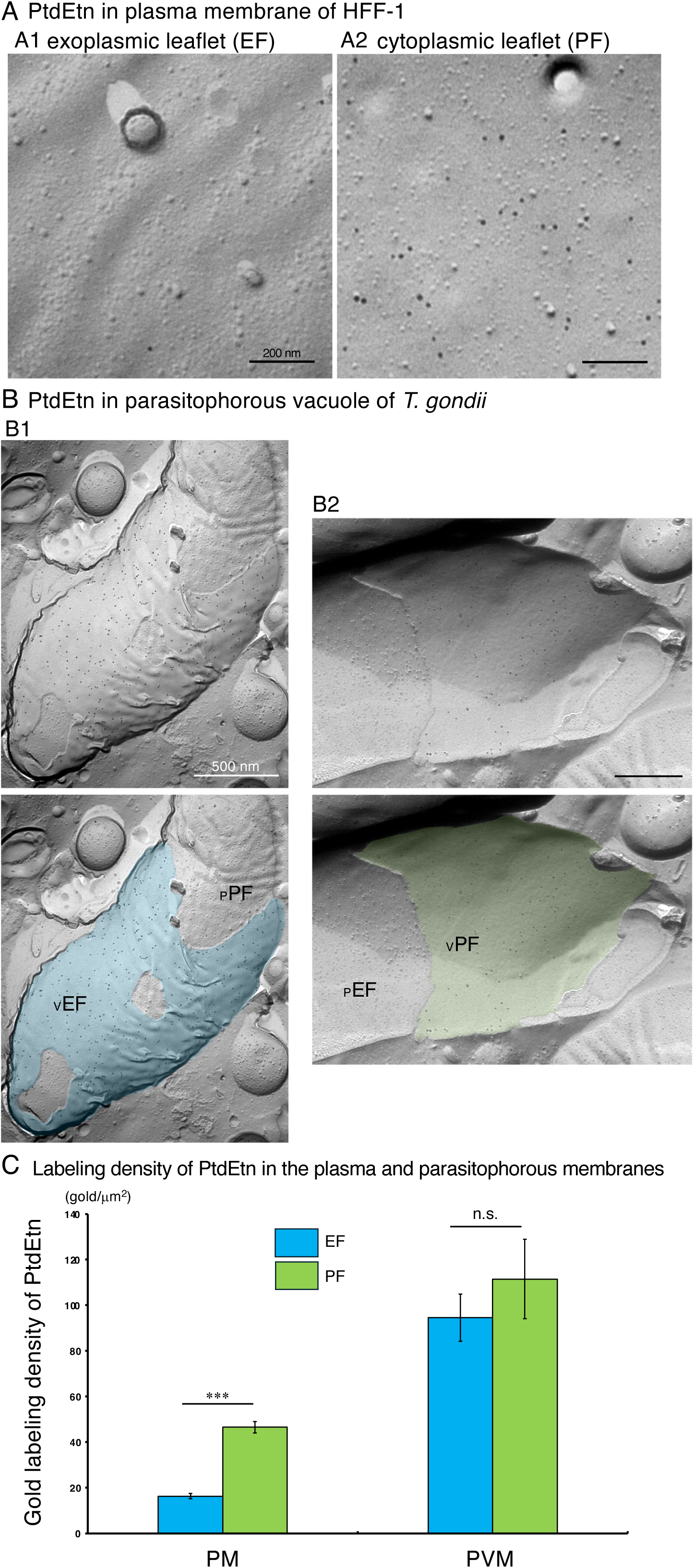
Distribution of PtdEtn in the HFF-1 PM and *T. gondii* PVM. Replicas were labeled with biotin-duramycin, which specifically binds to PtdEtn. (A) Replica images showing asymmetric localization of PtdEtn in the EF (A1) and PF (A2) of the HFF-1 PM (PF > EF). (B) Symmetric distribution of PtdEtn across the EF (blue) and PF (green) of the *T. gondii* PVM. _V_PF, _V_EF, _P_PF, and _P_EF indicate the P-face and E-face of the PVM and PM of *T. gondii*, respectively. (C) Quantification of gold labeling densities of PtdEtn. Scale bars: 200 nm (A), 500 nm (B). n. s.: not significant, *t*-test, ***, *p* < 0.001

### 3-4. Symmetrical localization of GM3 ganglioside in the PVM of T. gondii-infected cells

We next analyzed GM3 ganglioside, a glycosphingolipid known to reside exclusively in the EF of the PM. In uninfected HFF-1 cells, GM3 labeling was detected solely on the EF (Fig. 4A), consistent with its canonical asymmetric distribution. In contrast, the PVM of *T. gondii*-infected cells exhibited GM3 labeling on both the EF and PF (Fig. 4B), demonstrating that glycosphingolipid asymmetry is also lost upon PVM formation. Quantitative data (Fig. 4C) confirmed equivalent GM3 densities on both leaflets of the PVM.

**Fig. 4.**
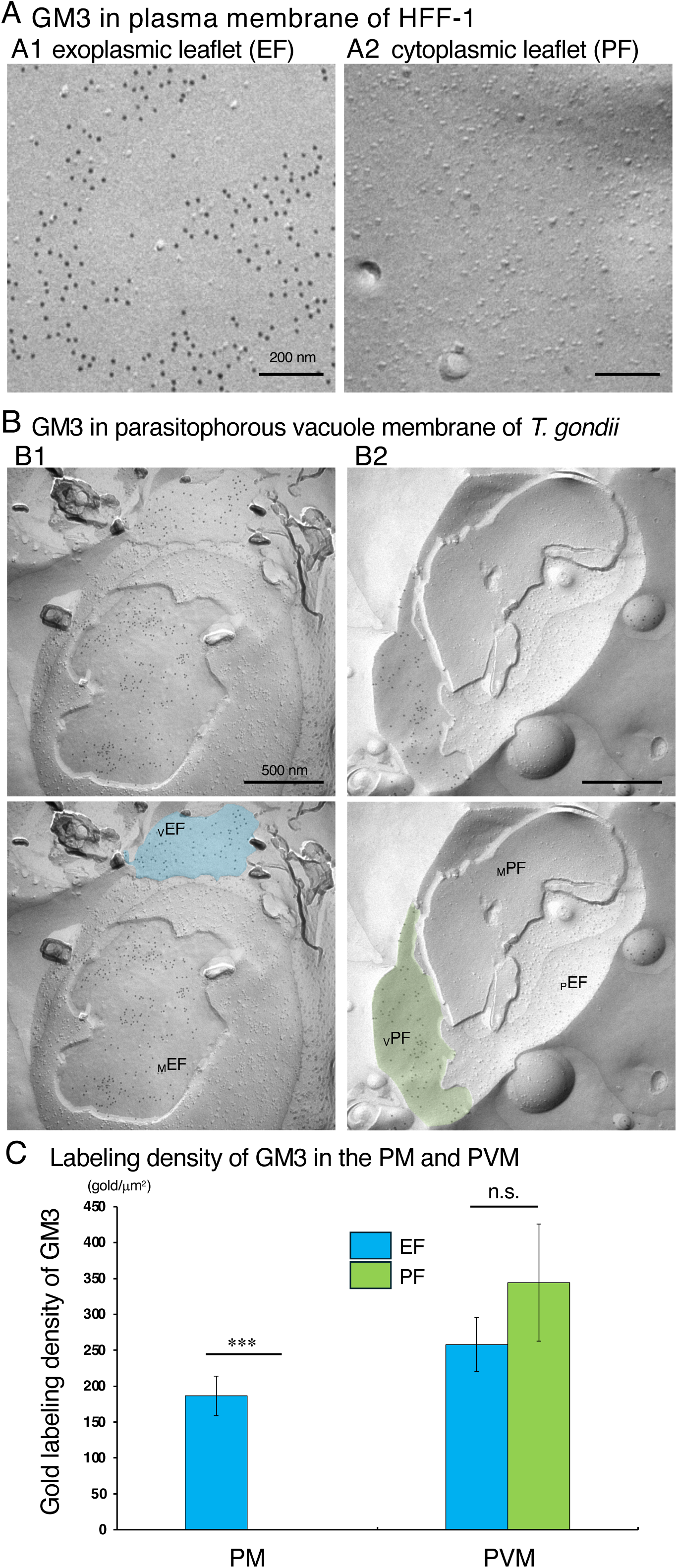
Distribution of GM3 ganglioside in the HFF-1 plasma membrane and *T. gondii* PVM. (A) Replica image showing GM3 exclusively in the EF of the HFF-1 PM. (B) Replica image showing symmetric GM3 distribution across both leaflets of the *T. gondii* PVM. _V_PF, _V_EF, _P_PF, and _P_EF indicate the P-face and E-face of the PVM and PM of *T. gondii*, respectively. (C) Quantitative comparison of GM3 labeling densities in EF and PF. Scale bars: 200 nm (A), 500 nm (B). n. s.: not significant, *t*-test, ***, *p* < 0.001

### 3-5. Symmetrical lipid distribution in the PVM of P. falciparum-infected erythrocytes

We extended our analysis to *P. falciparum*–infected human erythrocytes. Using the QF-FRL method, the erythrocyte PM exhibited characterstic membrane protrusions known as knobs. Using the freeze-fracture electron microscopy technique, these knobs were observed on both EF (arrows in Fig. 5A2, left) and PF (arrows in Fig. 5A2, right) when viewed from the hydrophobic interface. We found that in both uninfected (Fig. 5Aa) and *P. falciparum*–infected (Fig. 5A2) erythrocyte, PtdSer labelings were restricted to the PF of the PM. Similarly, PtdEtn labeling in erythrocyte PMs was predominantly localized to the PF, consistent with its inner leaflet enrichment (Fig. 6A). In contrast, GM3 labeling was confined to the EF of the PM and was absent from the PF in both uninfected and infected erythrocytes (Fig. 7A). In *P. falciparum*–infected erythrocytes, the PVM displayed symmetric PtdSer labeling across both leaflets (Fig. 5B). Furthermore, PtdEtn (Fig. 6B) and GM3 (Fig. 7B) labelings in *P. falciparum*–infected cells also appeared on both the EF and PF of the PVM. These findings are consistent with our observations in *T. gondii*, supporting the notion that lipid asymmetry is generally lost during PV formation in apicomplexan parasites.

**Fig. 5.**
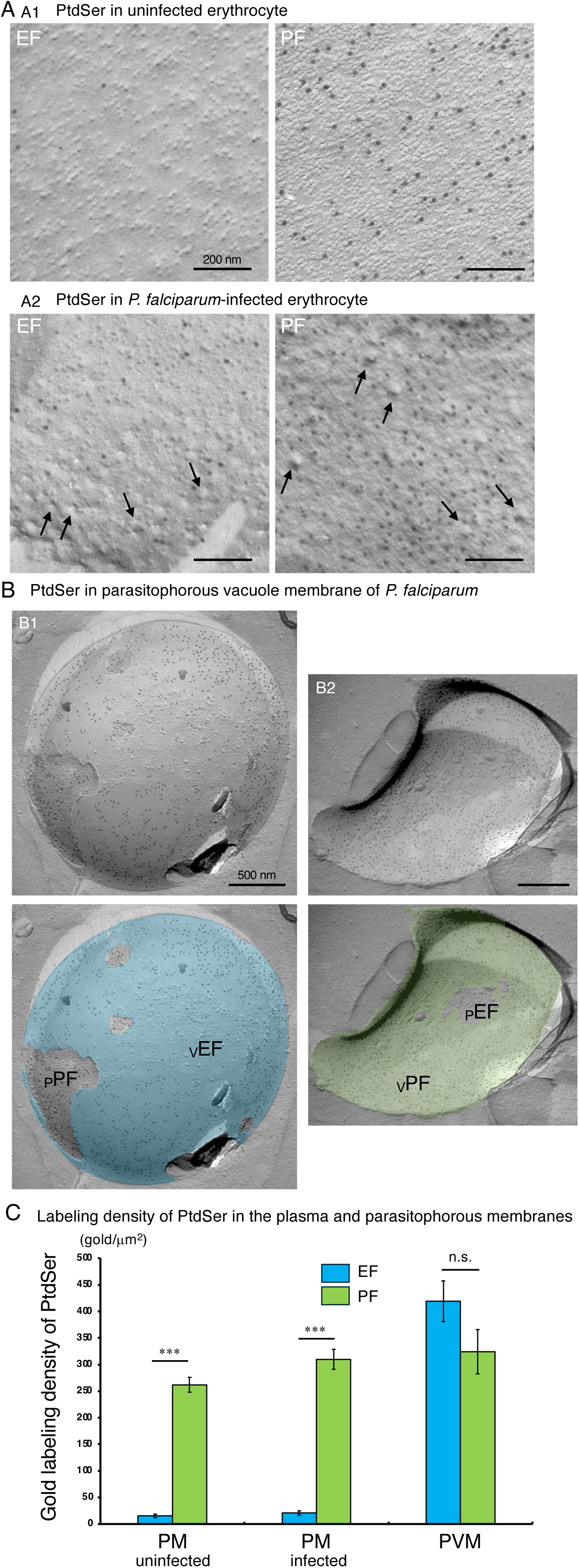
Distribution of PtdSer in the PM and PVM of *Plasmodium falciparum*–infected erythrocytes. (A) Replica images showing asymmetric distribution of PtdSer in both the uninfected and *P. falciparum*-infected erythrocyte PM (PF > EF). In *P. falciparum*-infected erythrocytes (A2), knob structures were clearly visible as indentations on the EF (arrows, left panel) and protrusions on the PF (arrows, right panel). (B) Replica images of the *P. falciparum* PVM showing symmetric distribution of PtdSer. _V_PF, _V_EF, _P_PF, and _P_EF indicate the P-face and E-face of the PVM and PM of *P. falciparum*-infected erythrocyte, respectively. (C) Quantitative analysis of PtdSer labeling in EF and PF of the PM and PVM. Scale bars: 200 nm (A), 500 nm (B). n. s.: not significant, *t*-test, ***, *p* < 0.001

**Fig. 6.**
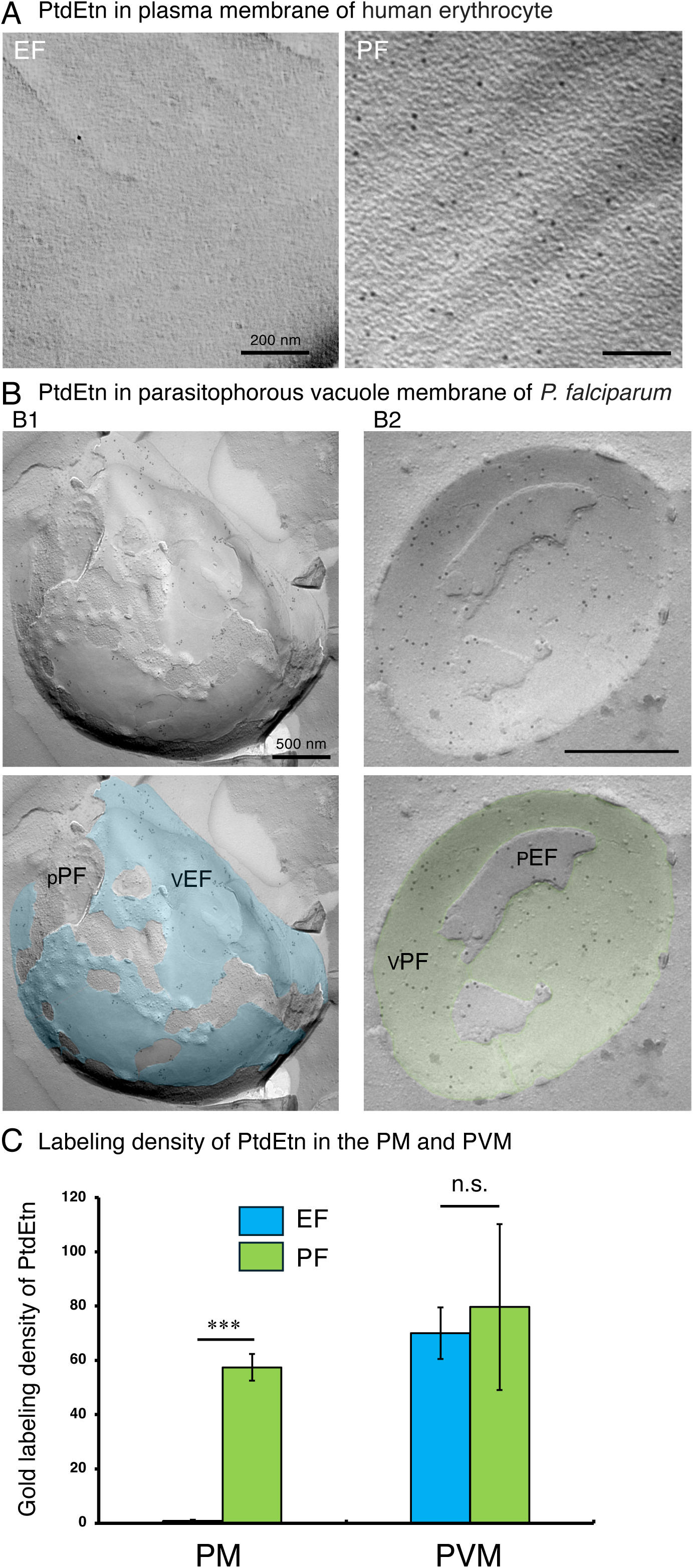
Distribution of PtdEtn in the plasma membrane of uninfected erythrocyte and PVM of *Plasmodium falciparum*–infected erythrocytes. (A) Replica images showing asymmetric distribution of PtdEtn in the erythrocyte PM (PF > EF). (B) Replica images of the *P. falciparum* PVM showing symmetric distribution of PtdEtn. _V_PF, _V_EF, _P_PF, and _P_EF indicate the P-face and E-face of the PVM and PM of *P. falciparum*-infected erythrocyte, respectively. (C) Quantitative analysis of PtdEtn labeling in EF and PF. Scale bars: 200 nm (A), 500 nm (B). n. s.: not significant, *t*-test, ***, *p* < 0.001

**Fig. 7.**
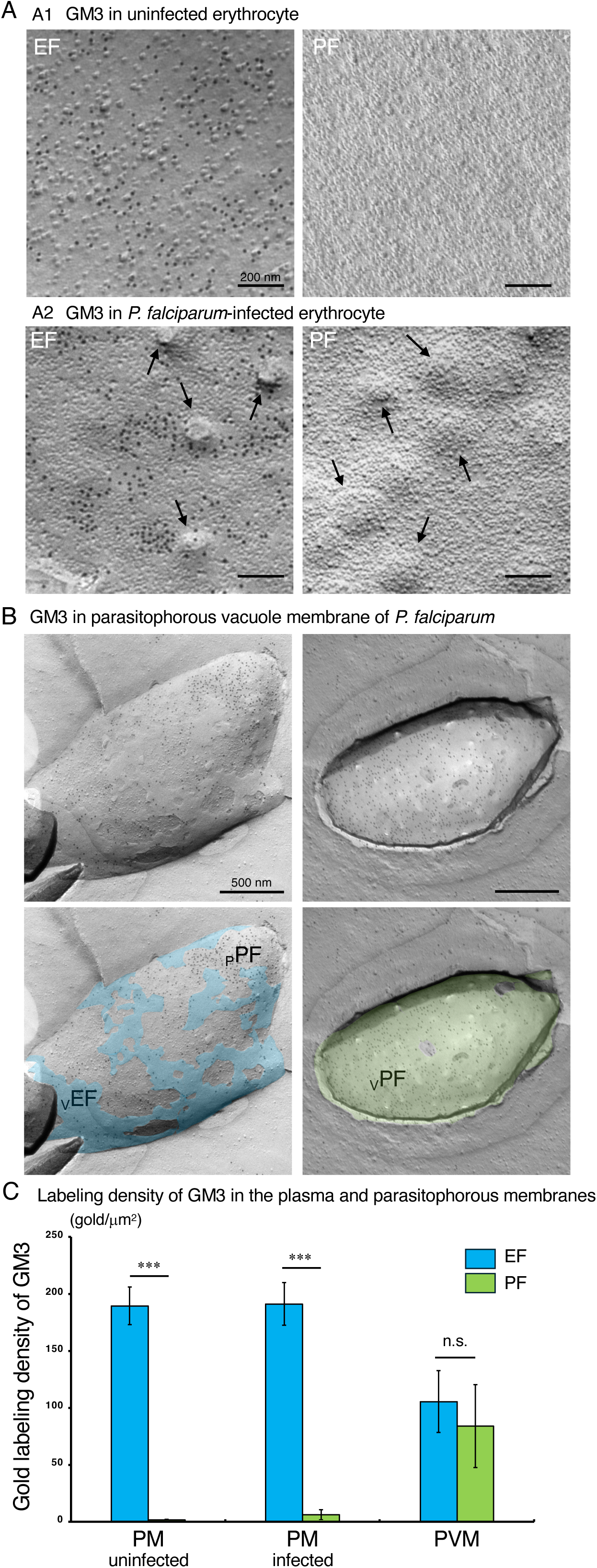
Distribution of GM3 in the PM of host erythrocyte and PVM of *P. falciparum*. (A) Replica image showing asymmetric distribution of GM3 in the erythrocyte PM (EF > PF). In *P. falciparum*-infected erythrocytes (A2), knob structures were clearly visible as indentations on the EF (arrows, left panel) and protrusions on the PF (arrows, right panel). (B) Replica image showing symmetric GM3 labeling across EF (blue) and PF (green) of the PVM. _V_PF, _V_EF, _P_PF, and _P_EF indicate the P-face and E-face of the PVM and PM of *P. falciparum*-infected erythrocyte, respectively. (C) Quantitative comparison of GM3 densities in the uninfected and *P. falciparum*-infected erythrocyte PM, and the *P. falciparum* PVM. Scale bar: 200 nm (A), 500 nm (B). n. s.: not significant, *t*-test, ***, *p* < 0.001

### 3-6. Distribution of IMPs in host and parasitophorous membranes

Because integral membrane-spanning proteins are often visualized as IMPs in freeze-fracture replicas [26–28], we next compared their density between host PMs and PVMs. As expected, the PM of uninfected HFF-1 cells (Fig. 8A1, B) and human erythrocyte (Fig. 8C1, D) exhibited a dense array of IMPs on the PF. In contrast, both the EF and PF of the PVMs in *T. gondii*-(Fig. 8A2, B) and *P. falciparum*-(Fig. 8C2, D) infected cells showed markedly reduced IMP densities. Quantitative analysis confirmed that IMPs were significantly low in the PVM (Fig. 8C, D). These data indicate that the PVM contains few integral membrane-spanning proteins, consistent with prior reports that host transmembrane proteins are largely excluded from the nascent PV during invasion [4, 5]. Interestingly, the IMP density on the PF was significantly higher than that on the EF in the PVM of *T. gondii*-infected cells, whereas no significant difference between the PF and EF was observed in the PVM of *P. falciparum*-infected erythrocytes (Fig. 8C, D).

**Fig. 8.**
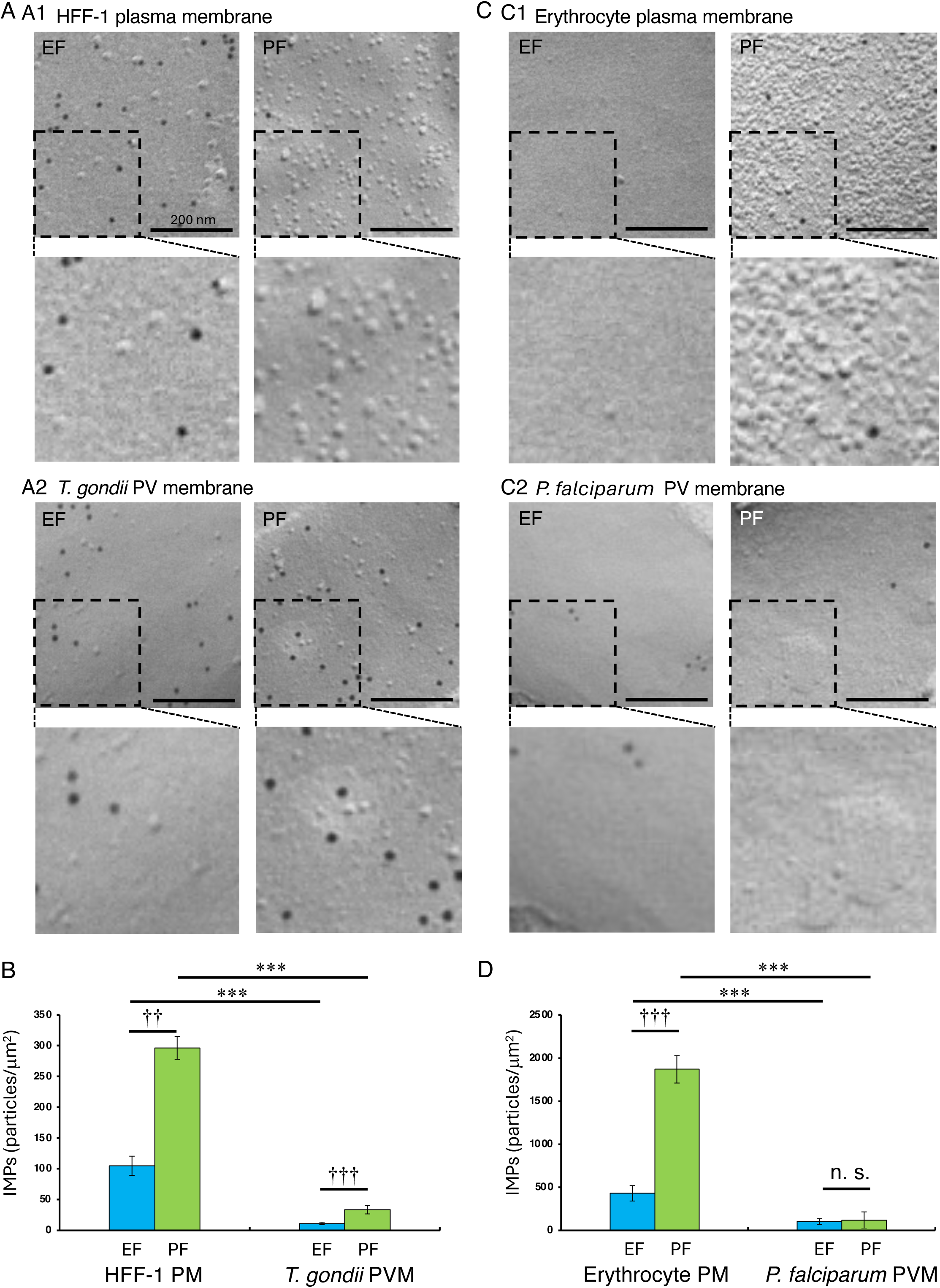
Analysis of intramembrane particle (IMP) distribution in PM of HFF-1 cell and human erythrocyte, and PVM of *T. gondii* and *P. falciparum*. Representative freeze-fracture replicas showing sparse IMPs in PM of HFF-1 cell and human erythrocyte, and PVM of *T. gondii* (A) and *P. falciparum* (C). Enlarged insets (lower panels) are included to better visualize the intramembranous particles and immunogold labeling. Histograms summarize IMP density on EF and PF leaflets (mean ± SD) of PM and PVM (B and D). Scale bars: 200 nm. *t*-test, ***: *p* < 0.001, ††: *p* < 0.005, †††: *p* < 0.001, n.s.: not significant.

**Fig. 9.**
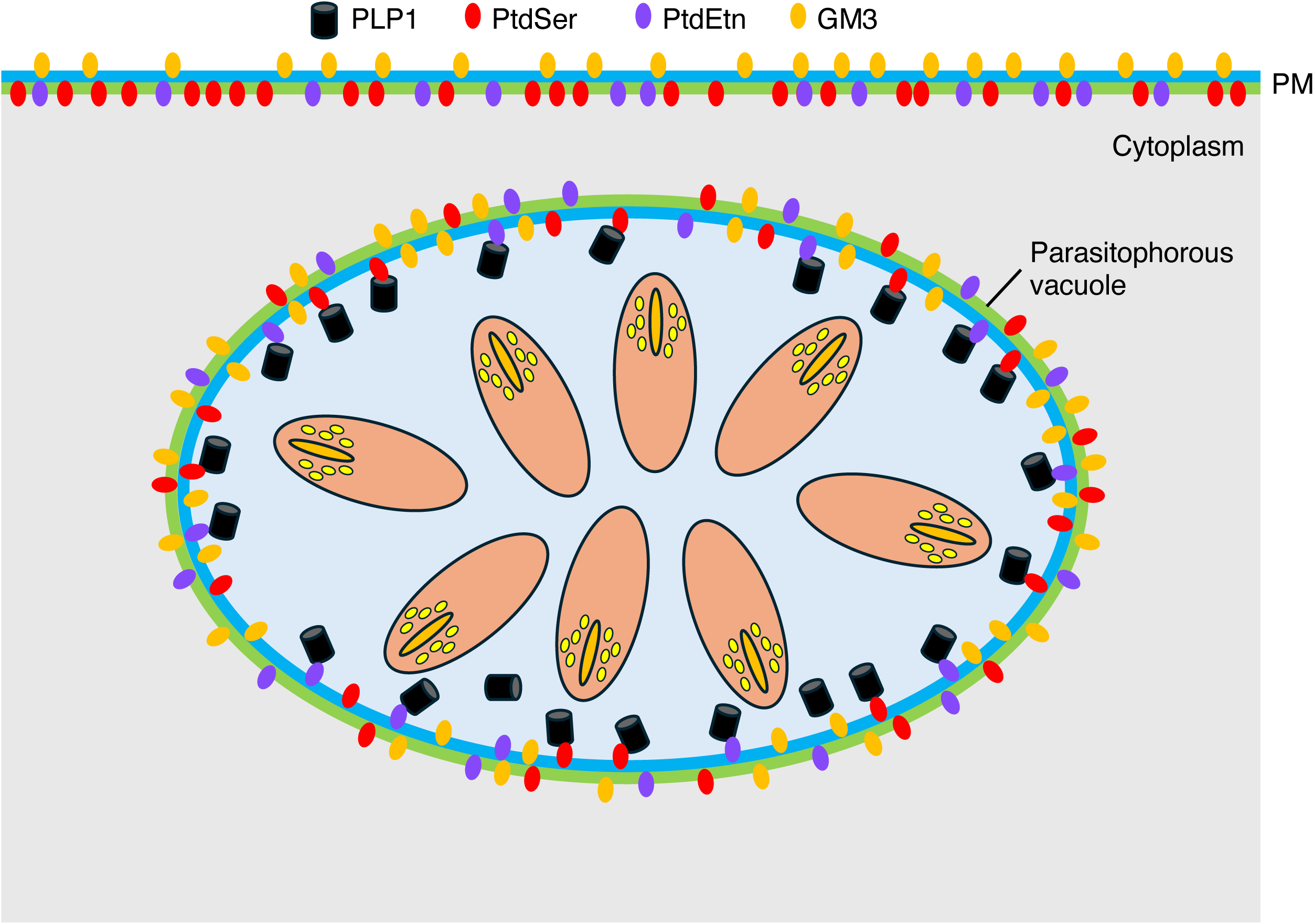
Schematic representation of PtdSer, PtdEtn, and GM3 distribution in the plasma membrane of host cells and the PVM in *T. gondii* and *P. falciparum*. In parasite-infected host cells, PtdSer (red) and PtdEtn (purple) are predominantly confined to the cytoplasmic leaflet (green) of the host plasma membrane (PL). However, in the PVM—derived from host plasma membrane during invasion—both PtdSer and PtdEtn are symmetrically distributed across the cytoplasmic (green) and luminal (blue) leaflets. This symmetric localization enables perforin-like protein 1 (PLP1, black), secreted by the parasite into the PV lumen, to interact with the inner (luminal) leaflet of the PVM via PtdSer and PtdEtn. This interaction facilitates membrane insertion and subsequent pore formation by PLP1, which is critical for parasite egress from the host cell.

## 4. Discussion

In this study, we used the QF-FRL technique to examine the transbilayer distribution of PtdSer, PtdEtn, and GM3 ganglioside in the PVM of *T. gondii* and *P. falciparum*. Our results revealed that all three lipids are symmetrically distributed across the EF and PF leaflets of the PVM. This strikingly contrasts with their canonical asymmetric distribution in host PMs, in which PtdSer and PtdEtn are enriched in the PF, while GM3 is restricted to the EF.

The loss of lipid asymmetry in the PVM suggests that profound membrane remodeling occurs during infection. This remodeling likely reflects a combination of physical and biochemical processes that act to equilibrate lipid distributions between the two leaflets of the membrane. Although spontaneous lipid flip-flop is generally slow under physiological conditions, the unique biophysical environment generated during parasite invasion—such as membrane curvature, tension, and local perturbation of lipid–protein interactions—may facilitate transbilayer movement. Alternatively, lipid translocases (scramblases or flippases) could be involved, although it remains unclear whether these enzymes are derived from the parasite or the host. Scramblases and flippases are not generally considered to be constitutively anchored to the membrane cytoskeleton [29–34], thus it is possible that these proteins are incorporated to the PVM. Identification of such proteins and clarification of their roles in PVM biogenesis and maintenance will be important subjects for future studies.

Our observation that the PVM contains very few IMPs provides additional insight into its unique architecture. Previous studies have shown that integral host membrane proteins, including CD44, β1-integrin, and Na⁺/K⁺-ATPase, are excluded from the PVM [4, 7]). In contrast, glycosylphosphatidylinositol (GPI)-anchored proteins can traverse the moving junction and remain associated with the PVM (Mordue et al. 1999). Our QF-FRL data confirm that IMPs—corresponding mainly to integral transmembrane proteins [26–28]—are largely absent from both the EF and PF of the PVM, consistent with the exclusion of most host transmembrane proteins. This finding raises an intriguing question regarding the apparent scarcity of parasite-derived integral membrane proteins, particularly dense granule (GRA) proteins. It is well established that numerous GRA proteins become integrated into the mature PVM [35, 36], many of which contain a single transmembrane domain [37]. Such single-pass proteins may not produce prominent intramembrane particles in freeze-fracture replicas, in contrast to multi-pass proteins or large protein complexes that create more prominent IMPs, such as the MYR translocon in *T. gondii* and PTEX in *Plasmodium* [27] [38] [39]. Therefore, the low IMP density observed in our analyses of PVs collected 2–3 days post-infection does not necessarily indicate the absence of GRA proteins but likely reflects a limitation of the freeze-fracture technique in detecting small or dispersed single-pass transmembrane proteins. As detailed in the Methods, the analyses were performed using *T. gondii*-infected HFF-1 cells at 2–3 days post-infection, when PVM remodeling and GRA protein incorporation are largely complete. The presence of PtdSer and PtdEtn on the luminal (parasite-facing) leaflet of the PVM may have important functional implications for parasite egress. *T. gondii* secretes perforin-like proteins (TgPLP1) that are essential for efficient egress [40]. The C-terminal domain (CTD) of TgPLP1 preferentially binds to membranes enriched in PtdSer and PtdEtn [41]. The exposure of these lipids on the luminal leaflet of the PVM would therefore facilitate PLP1 binding and pore formation, allowing the parasites to rupture the PVM during egress (Fig. 8). The symmetrical distribution of these aminophospholipids may thus represent a preparatory step for egress, ensuring that the PVM provides appropriate binding substrates for PLP1-mediated disruption.

We found higher IMP density on the PF than EF in the *T. gondii* PVM, but no PF–EF difference in the *P. falciparum* PVM (Fig. 8). This may reflect the fact that *T. gondii* resides metabolically active nucleated cells and recruits host mitochondria and various host molecules to the PVM [42, 43]. In contrast, mature erythrocytes infected by *P. falciparum* lack a nucleus and intracellular organelles, and there have been no reports of host molecules being recruited to the PVM.

In summary, our findings reveal that the PVM of both *T. gondii* and *P. falciparum* loses canonical lipid asymmetry and instead exhibits a symmetric organization of phospholipids and glycosphingolipids. This unique feature likely results from membrane remodeling during invasion and may have significant implications for parasite survival, immune evasion, and egress. Future studies identifying the molecular machinery responsible for this lipid redistribution—whether parasite-encoded or host-derived—will be essential for understanding the biogenesis and function of this distinctive membrane. Furthermore, an interesting subject for future studies will be to determine whether the loss of lipid asymmetry is established immediately after parasite invasion or is gradually acquired during maturation of the parasitophorous vacuole membrane.

## Data availability

All data generated or analyzed during this study are included in this published article (and its Supplementary information files).

